# Utilising hippocampal neuronal calcium activity in mouse CA1 for a multimodal optical brain-computer interface

**DOI:** 10.1101/2022.04.26.489497

**Authors:** Dechuan Sun, Forough Habibollahi Saatlou, Yang Yu, Ranjith Rajasekharan Unnithan, Chris French

**Affiliations:** Department of Medicine, The University of Melbourne, Victoria, Australia; Department of Electrical and Electronic Engineering, The University of Melbourne, Victoria, Australia

## Abstract

The hippocampus has been proposed to integrate information from multiple sensory modalities, supporting a comprehensive “cognitive map” for both spatial and non-spatial information. Previous studies have demonstrated decoding of hippocampal spatial information in real time by recording neuronal action potentials with electrodes. However, decoding of hippocampal non-spatial information robustly in real-time has not been previously shown. Here, we utilise the advantages of widefield optical calcium imaging to construct an optical brain-computer interface (BCI) driven by calcium activity of large neuronal ensembles (∼600 neurons) to decode spatial, visual and auditory information effectively in real time. We developed a high speed end-to-end analysis workflow with advanced machine learning techniques for decoding. This methodology achieves high decoding accuracy and provides a “cognitive translation” approach that may be applied to both research and clinical applications to allow direct neural communication with animals and patients with impairment of function.

## 1. Introduction

The hippocampus, embedded deep within the temporal lobe, connects with many other brain structures directly or indirectly and plays important roles in memory and cognition. It has been demonstrated to support a “cognitive map”, providing an environment-centric spatial memory system (O’keefe and Nadel, 1978; Spiers et al., 2020). Apart from encoding spatial information, previous studies have revealed that the hippocampal neuronal network also encodes non-spatial information such as visual (Acharya et al., 2016; Liu et al., 2018), auditory (Moita et al., 2003; Itskov et al., 2012; Xiao et al., 2018), odour (Komorowski et al., 2009; Taxidis et al.,2020), gustatory (Ho et al., 2011), and tactile information (Pereira et al., 2007; Gener et al., 2013). These intriguing observations indicate a more abstract and comprehensive hippocampal cognitive map, generating a high-dimensional space for both spatial and non-spatial information. Accurately decoding the hippocampal cognitive map in real time would significantly enhance direct neural communication (“cognitive decoding”) in both experimental and clinical scenarios.

Hippocampal spatial information has previously been decoded in real time using electrophysiological signals (Guger et al., 2011; Sodkomkham et al.,2016; Ciliberti et al. 2018; Hu et al., 2018). However, whether it is possible to decode hippocampal non-spatial information in real time has not yet been studied, possibly due to limitations on the number of electrophysiological recording channels, complicated and time-consuming analyses pipelines, and relatively low sensitivity of hippocampal neurons to non-spatial information. Additionally, the electrodes often shift and the signal tends to attenuate over time.

Here, we describe an optical brain-computer interface (OBCI) based on a single-photon imaging technique (“miniscope”, Ghosh et al., 2011) to decode hippocampal spatial, visual, and auditory information in three experiments. First, spatial position was assessed with the animals traversing a linear track. In the second and third experiments, the animals were placed into a chamber and passively exposed to light stimuli or pure sinusoidal tones centred at three different frequencies. The OBCI detects activity of very large neuronal ensembles and provides stable calcium activity for very long time periods. Instead of implementing traditional BCI analysis pipelines to detect action potentials, we developed an end-to-end workflow using the raw calcium activity data together with machine learning models for decoding without inferring neuron spikes. To the best of our knowledge, this is a unique methodology in OBCI implementation. Our analysis pipelines offer considerable promise for interpreting this high bandwidth multimodal dataset. We achieved a decoding error of 9.33 ± 0.67 cm/frame (mean ± standard error mean) in position reconstruction experiments, a decoding error ratio of 3.36% ± 1.47% in visual stimuli identification experiments, and 17.83% ± 2.32% in auditory stimuli identification experiments. This study presents for the first time a proof of concept for decoding the hippocampal cognitive map in real time for “cognitive decoding” using an end-to-end optical brain computer interface.

## 2. Materials and Methods

All procedures were approved by the Florey Animal Ethics Committee (No. 18-008UM), subject to the restrictions contained in the Australian Code for the Care and Use of Animals for Scientific Purposes, 8th Edition.

### 2.1 Animals and surgery

C57BL/6J mice (8 weeks old) were used in the experiment and underwent two stereotaxic surgeries under anaesthesia (isoflurane: 3%–5% induction, 1.5% maintenance). Mice were first unilaterally injected with 500nl of pAAV.Syn.GCaMP6f.WPRE.SV40 virus (#100837-AAV1, AddGene) in dorsal CA1 (right hemisphere, 2.1 mm posterior to the bregma, 2.1 mm lateral to the midline, and 1.65mm ventral from the surface of skull) over a 15 min period. One week following the injection, two anchor screws were secured to the skull and a circular craniotomy 2mm in diameter was made next to the injection site (2.1 mm posterior to the bregma, 1.6 mm lateral to the midline). The cortex above the corpus callosum was aspirated using a 27-gauge blunt needle and a vacuum pump, and a 1.8mm diameter GRIN lens (#64-519, Edmund Optics) was implanted at a depth of 1.35mm from the surface of the skull. Cyanoacrylate glue and dental acrylic were used to fix the lens in place, and the lens was protected by silicone adhesive (Dragon Skin® Series). The mice were given analgesics (carprofen:5mg/kg; dexamethasone: 0.6mg/kg) and enrofloxacin water (1:150 dilution, Baytril®) to recover for seven days. The neuronal calcium activity was examined 4-5 weeks later. After finding the best field of view, a baseplate was cemented on the animal’s head and a plastic cap was locked into the baseplate to protect the lens (**Fig 1a-c**).

**Figure 1.**
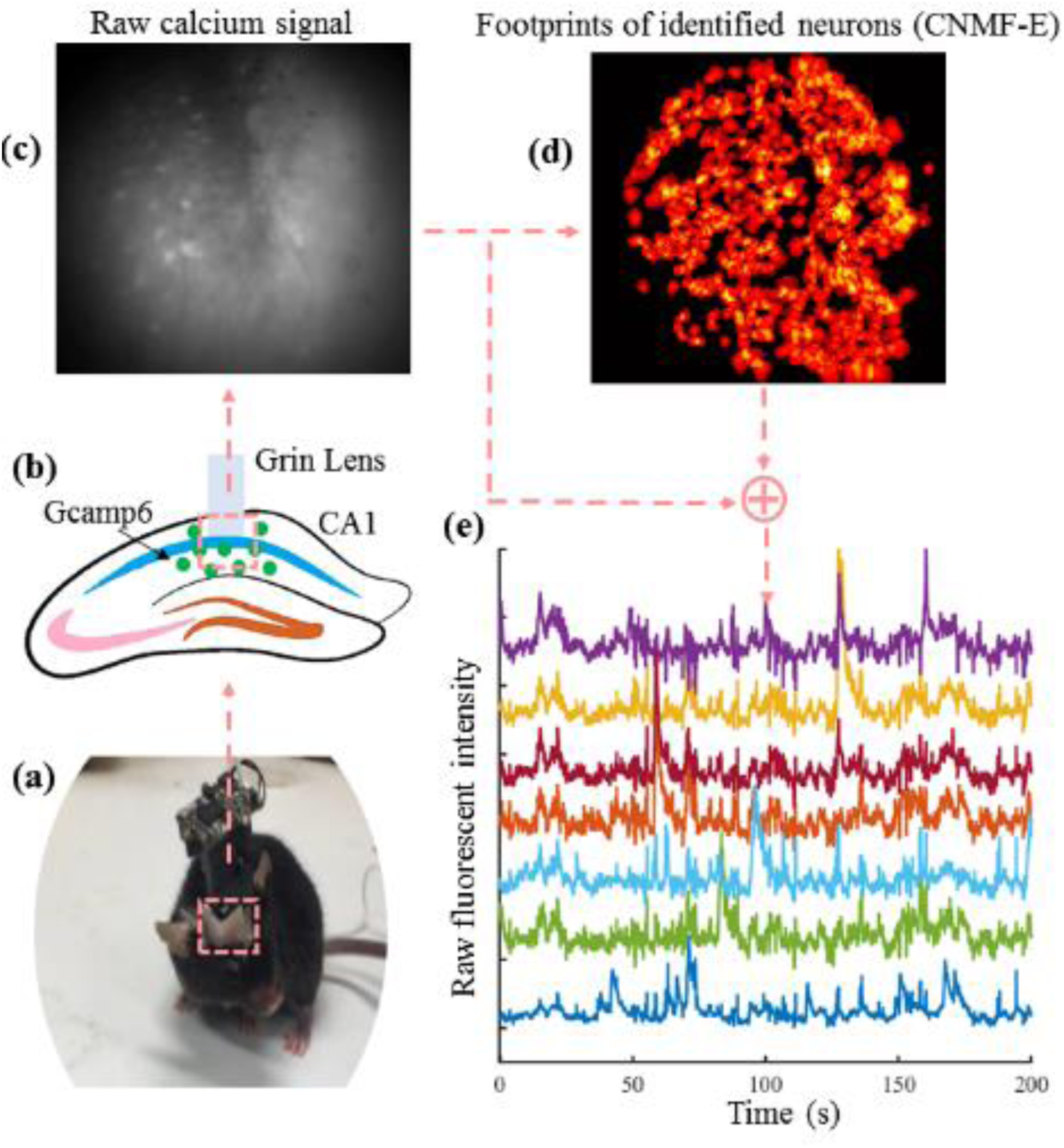
In vivo calcium signal recording pipelines. **(a)** An example of a mouse with a miniscope. **(b)** Diagram of Grin lens implanted in hippocampal CA1. **(c)** An example of the raw calcium signal. **(d)** An example of the spatial footprints of neurons identified with CNMF-E algorithm. **(e)** An example of the raw fluorescent intensity of detected neurons.

**Figure 2.**
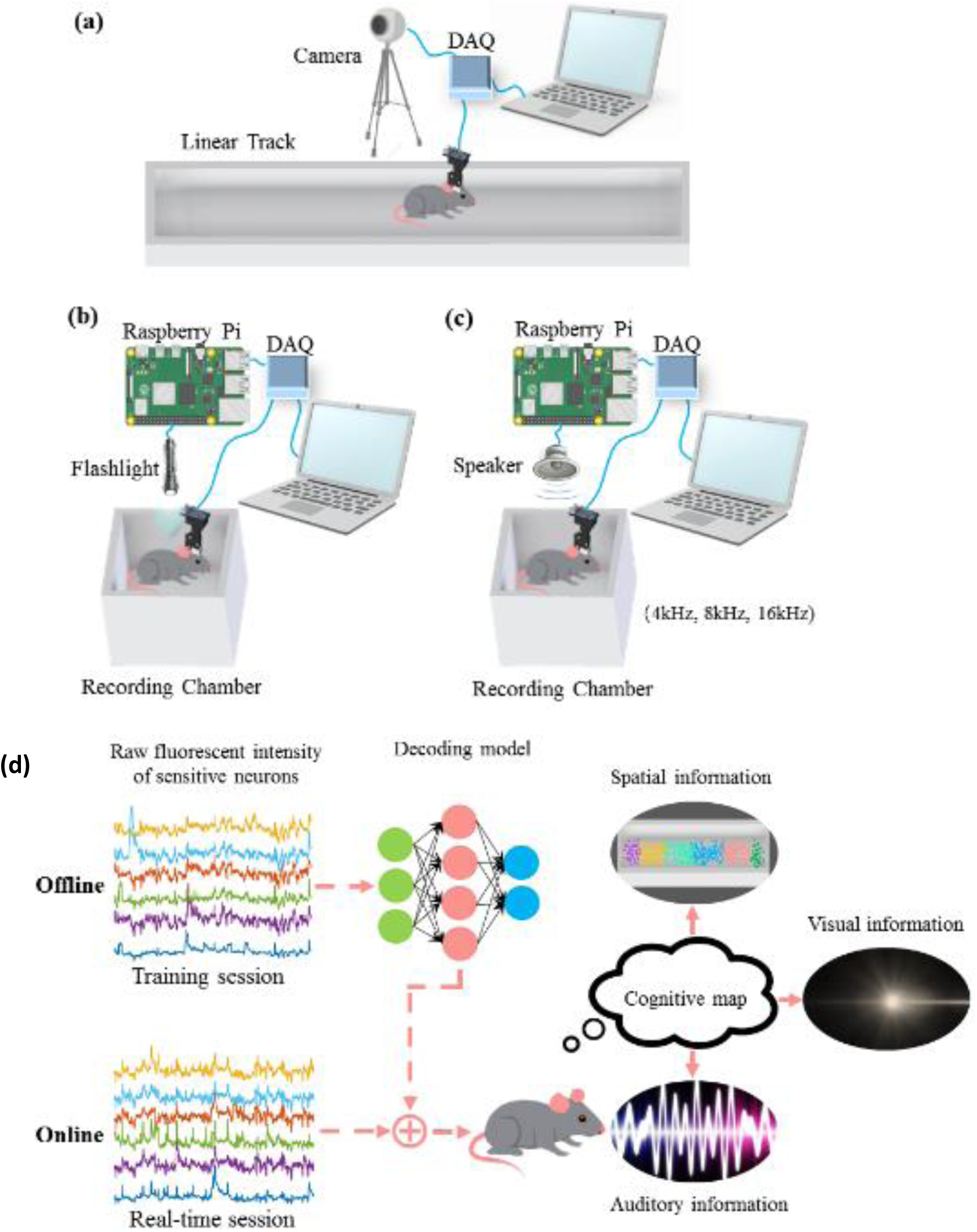
Experimental apparatus. The miniscope and data acquisition board (DAQ) were used to record the animal’s hippocampal calcium activity during experiments. **(a)** The mouse traversed a 1.6m linear track and a camera was used to track the animal’s location. **(b-c)** The animal was placed in a small recording chamber. The visual and auditory stimuli were controlled using a Raspberry Pi. **(b)** The flashlight was switched on and off alternately. **(c)** The speaker was activated and deactivated alternately, playing sinusoidal tones centred at 4kHz, 8kHz, or 16kHz randomly. **(d)** Hippocampal activity decoding pipelines. In the training session, the raw fluorescent intensity of sensitive neurons was used to construct the decoding model. This model was then deployed in the real-time session to decode the spatial, visual, and auditory information.

### 2.2 Information content and sensitive neurons

Information content is a measure to quantify the precision level of neuronal coding; a larger value indicates more precise coding. In the position reconstruction experiment, the definition of information content is similar to that described in Ravassard et al. (2013) and Rubin et al. (2015), but the measurement “neuronal firing rate” is changed to “neuronal fluorescent intensity” to adapt to our recording technique,

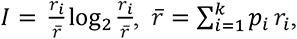

where *I* represents the information content, *i* is the spatial bin index, *k* is the number of spatial bins, *p_i_* is the probability of occupancy of the *i*^th^ bin, *r_i_* is the average fluorescent intensity in the *i*^th^ bin and *r* is the overall mean fluorescent intensity. We next shuffled the animal’s location, and the information content was recomputed for each shuffle. This step was repeated 1000 times, and a neuron was marked as a location sensitive neuron if its information content in the unshuffled trial exceeded 95% of the shuffled trials.

In visual and auditory stimuli experiments, the definition of the information content was the same, but the data binning was implemented in the temporal domain. There were two states (light or dark) in the visual experiment, and three states (three different frequencies) in the auditory experiment. We used the same shuffling method to detect light and sound sensitive neurons.

### 2.3 Calcium activity decoder

We tested and compared the performance of several decoders to decode the neural signals including a Gaussian naïve Bayes (GNB) decoder, a support vector machine (SVM) decoder, a multilayer perceptron (MLP) neural network model, a long short-term memory (LSTM) neural network model, and a convolutional neural network (CNN) model. The decoders were constructed and implemented on the Python platform using the scikit-learn tool kit (Pedregosa et al., 2011) and tensorflow library (Abadi et al., 2016).

The GNB decoder is a type of probabilistic-based prediction algorithm based on Bayes’ theorem. In the position reconstruction experiment, raw fluorescent intensities of location sensitive neurons were first normalized to remove the mean and scaled to unit variance. Given a sequence of fluorescent intensities from location sensitive neurons in each frame, the estimated position *y* was defined as,

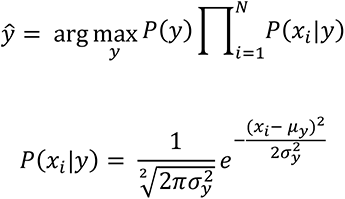

where *P*(*y*) is the probability of occupancy of bin *y*, *x_i_* is the normalized fluorescent intensity of the *i*^th^ neuron, and *μ_y_* and *σ_y_* represent the mean and standard deviation of the normalized fluorescent intensity of the *i*^th^ neuron at bin *y*, respectively. In the visual stimuli experiment, *y* represents either light-bin or dark-bin. In the auditory stimuli experiment, *y* represents the bins with different stimulus frequencies.

The SVM decoder performs classification by constructing a set of hyperplanes that maximizes the margins between different classes. The SVM model was trained and constructed using the normalized input data with a non-linear radial basis function kernel. The kernel width, gamma, was set to be the reciprocal of the number of input features. A cost parameter “C” was optimized by applying a grid search technique with five-fold cross-validation.

An MLP is an artificial neural network that is commonly used in solving the problems of prediction and classification. It shows excellent performance when the input data is not linearly separable. The MLP neural network constructed in our experiments contained one input layer, two hidden layers activated by a rectified linear unit (ReLU) function, and one output layer with softmax activation. The input data was normalized and fed into the model. We used an Adam optimizer and a categorical cross-entropy loss function to compile the model. The batch size was set to be 32 and all the other hyperparameters including the number of nodes in hidden layers, learning rate, the number of epochs were optimized by implementing a grid search method using five-fold cross-validation.

An LSTM model is a type of recurrent neural network that can use internal memory to process sequences of data with variable length and has shown notable success in time series forecasting. The LSTM model was implemented to reconstruct the animal’s running trajectory in our experiments. The model consisted of one input layer, two fully connected LSTM layers activated by a ReLU function, a dropout layer (dropout rate: 0.2) that prevented overfitting, and one output layer with a softmax activation function. In the position reconstruction experiment, the output of the model was marked as the animal’s current location bin, and we tested different lengths of normalized time-series data to build and train the model. The same method as the MLP neural network was used to compile and tune the hyperparameters of the model.

A CNN is a class of artificial neural network that has been frequently implemented in image processing, but also shows good performance for time series data. It is designed to detect spatial hierarchies of features in the input data. We tested a CNN model to decode the hippocampal activity in auditory stimuli experiments. The inputs were the time series of raw fluorescent intensities from sensitive neurons. The model contained two convolutional layers with ReLU activations, with a max-pooling layer added after each convolutional layer for dimensionality reduction. A dropout layer (dropout rate: 0.2) was then concatenated to prevent overfitting. Finally, a fully connected layer with a softmax activation function was added to output the probability distribution for each class. A grid search method was performed to determine the hyperparameters that yielded the highest decoding accuracy.

### 2.4 Kalman filter

In the position reconstruction experiment, we used a Kalman filter to reduce the decoding noise in the outputs of the decoder. The Kalman filter is one of the most widely used methods for position tracking and estimation. To estimate the state *x̂_t_* [position and velocity] of the animal at time *t*, the estimation processes are defined as:

#### Time update

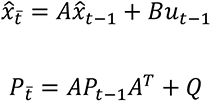

#### Measurement update

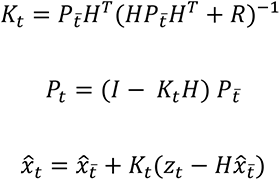

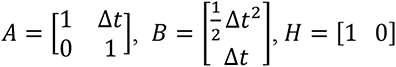

where *x̂_t̄_* is the prior estimation of the state, *P_t̄_* is the prior transition covariance, *K_t_* is the Kalman gain, *P_t_* is the updated transition covariance, *x̂_t_* is the updated state, *u_t-1_* is the acceleration, *Q* is the transition covariance and *R* is the observation covariance. The sampling frequency is 30 fps, so Δ*t* equals 1/30. The initial values of the state were set to be zero and the transition covariance matrix was set to be identity. The values of *Q* and *R* were set to be 0.0001 and 1, respectively.

### 2.5 Experimental procedures and analysis method

The experiment contained two sessions: (1) a training session and (2) a real-time session. The training session was implemented offline to detect the location of sensitive neurons from the raw calcium images and to construct the decoding model. In the real-time session, we used the decoding model constructed in the training session to decode the neuronal activity directly.

#### 2.6.1 Real-time position reconstruction

Before the experiment, the mice were kept on a dietary restriction and their body weights were maintained at 85% of free-feeding body weights. We trained the mice with the miniscope attached to traverse a 1.6m linear track for food rewards. The mice were required to complete 12 trials of traversing each day for one week.

##### Training session

On the day of the training session experiment, we first recorded the hippocampal calcium activity when the mouse traversed the linear track for 12 trials. The sampling frequency of the miniscope was set to 30 fps and the animal’s position was tracked using a video camera mounted above the linear track that was synchronized with the miniscope recording system. After recording, the translational frame shifting was corrected using a cross-correlation-based image registration algorithm (Guizar-Sicairos et al., 2008). The spatial footprints of neurons in the field of view were detected by implementing a constrained nonnegative matrix factorization for endoscopic recordings (CNMF-E) algorithm (Zhou et al., 2018; **Fig 1d**). The CNMF-E could effectively detect neurons in different layers, leading to the overlap of the neuronal footprints. To decrease interference from surrounding neurons, we used the centroid of each neuron and a small surrounding area to extract their calcium activity. For each neuron, the raw calcium activity was defined as the average fluorescent intensity of the centroid pixel and the surrounding eight pixels (**Fig 1e**). To quantify the spatial information content, the linear track was sectioned into 80 bins of 2cm and the time spent in each bin was measured. We then calculated the spatial information content of each neuron and detected the location-sensitive neurons using a shuffling method. Next, the calcium activity of location-sensitive neurons was used to construct a decoding model to assess how well the calcium activity of neuronal ensembles predicted the animal’s location. We tested and compared the performance of a GNB decoder, an SVM decoder, an MLP neural network model, and an LSTM neural network model. Finally, a Kalman filter was used to reduce the decoding noise from the outputs of the decoder. The best decoder together with the Kalman filter was subsequently deployed in the real-time session.

##### Real-time session

In the real-time session, the mouse traversed the linear track for 12 trials. The position of the mouse was tracked by a video camera, but it was only used later to assess the decoding accuracy. The same image registration method was implemented to align the image and the raw calcium activity of position sensitive neurons detected in the training session was extracted and fed into the decoder to reconstruct the animal’s running trajectory in real time.

#### 2.6.2 Visual and auditory stimuli identification

We separately studied hippocampal activity evoked by visual stimuli and auditory stimuli. The mouse was put in a small opaque recording chamber and was exposed to either a flashlight or a speaker mounted above the chamber.

##### Training session

On the day of the training session experiment, we placed the mouse in the chamber 15 minutes before recording to acclimate. We used a Raspberry Pi 3 (Model B) board to control a flashlight or a speaker. In the visual stimuli experiment, the flashlight was turned on 2s and off 2s alternately150 times. In the auditory stimuli experiment, the speaker played three different frequency tones (4kHz, 8kHz, 16kHz) randomly 225 times with an activation-period of 2s and a mute-period of 3s. The data was analysed using the same procedure described in the position reconstruction experiment: (1) image registration, (2) neuron centroid detection, (3) raw calcium activity extraction, (4) sensitive neuron detection, and (5) decoder construction. We tested and compared the performance of a GNB decoder, an SVM decoder, and an MLP neural network model. In the auditory stimuli experiment, none of these three decoders showed outstanding performance (see Results), so we divided the data into several epochs for further analysis. The time length of the epoch was 5s, which included the speaker activation and deactivation periods. Finally, we constructed a CNN model to decode the epoch data.

##### Real-time session

In the real-time session, the mouse was exposed to light stimuli 150 times or sound stimuli 80 times. The raw calcium activity of stimuli sensitive neurons detected in the training session was extracted after image registration and was provided to the decoder. An MLP model or an SVM model was deployed in the visual stimuli experiment and a CNN model was deployed in the auditory stimuli experiment to do the subsequent real-time decoding.

### 2.6 Noise level and firing rate map similarity

The noise level was used to characterize the noise coupled in the calcium dynamics and the background noise. The raw calcium activity was first processed with a zero-phase infinite impulse response lowpass filter (1Hz cut-off frequency, filter order:20), and the signal noise level was defined as the difference between the raw calcium activity and the filtered activity. To compare the consistency or the similarity of the firing rate map between the training session and the real-time session, we measured the normalized cross-correlation (Lewis, 1995)

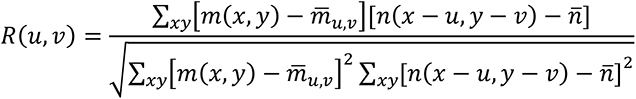

where *m* and *n* represent the firing rate maps in the training session and the real-time session, *x* and *y* represent the pixels in the maps, and *u* and *v* represent the pixel shift along different dimensions in each map, respectively.

## 3. Results

We constructed an optical brain-computer interface (OBCI) to decode mouse hippocampal calcium activity in real time in response to different sensory modality stimulation. We compared the performance of several decoders using the training data and separately measured real time decoding accuracy in a position reconstruction experiment (n=3 mice), a visual stimuli identification experiment (n=3 mice), and an auditory stimuli identification experiment (n=3 mice).

### 3.1 Position reconstruction

To investigate whether the raw hippocampal calcium activity could be decoded to reconstruct a mouse’s moving trajectory in real time, we tested three mice separately in a 1.6m linear track. A camera was synchronized with the miniscope recording system to capture the mouse’s running trajectory during the experiments. Each mouse first underwent a training session to construct the position reconstruction model. A large population of hippocampal neurons was observed in each mouse (n = 781, 478, 622, respectively). We measured the spatial information content of each neuron and detected the position sensitive neurons (n = 322, 320, 250, respectively) using a shuffling method (see Methods). An example of the place field map of sensitive neurons is shown in **Fig 3a**. The raw fluorescent intensities of sensitive neurons were used as “selected features” to train the position reconstruction model. We separately tested and compared the performance of a GNB decoder, an SVM decoder, an MLP neural network, and an LSTM neural network. To eliminate the reconstruction outliers, a Kalman filter was cascaded at the end (see Methods). Using five-fold cross-validation, we optimized the hyperparameters of each model and measured the reconstruction errors (summarized in **Supplementary Table 1**). All models could reconstruct the mice’s running trajectories accurately. The GNB decoder showed the highest decoding error (mean: 22.79 ± 3.42 cm/frame; median:16.00 ± 2.64 cm/frame), while the other three decoders achieved better performance with similar decoding errors (mean: ∼14 cm/frame; median: ∼11 cm/frame, **Fig 3b-c**). An example of the position reconstruction using the training data is shown in **Fig 3d**. Intriguingly, different firing patterns of sensitive neurons were observed and the running trajectory could be described in a high-dimensional neural state space (**Supplementary Fig 4.a,d**). Next, the decoding model showing the best performance was prepared to deploy in the real-time session.

**Figure 3.**
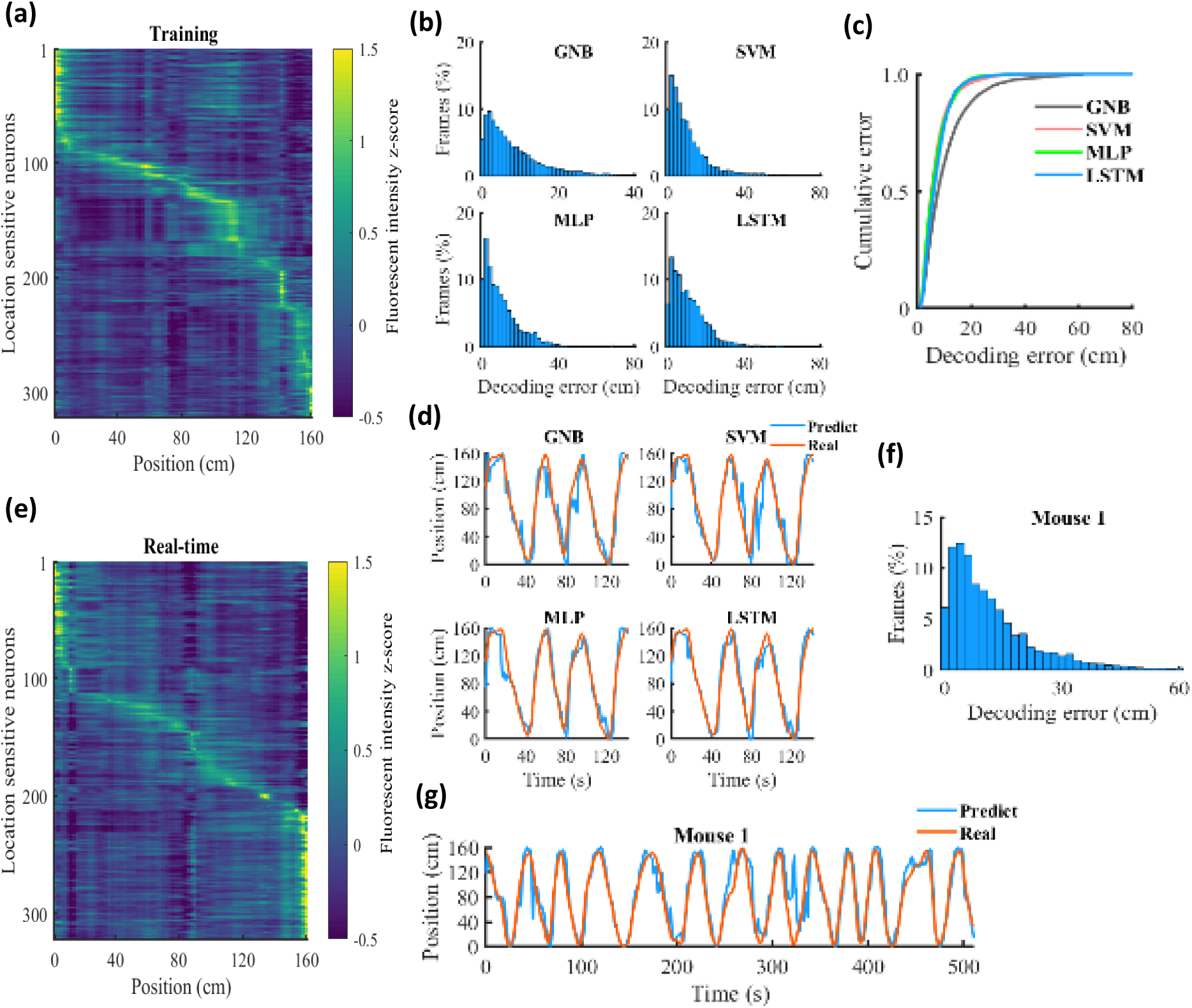
Position reconstruction experiment. **(a)** An example of the place field map of sensitive neurons in training sessions. The colour represents the fluorescent intensity z-score. **(b)** Decoding error histograms of five-fold cross-validation results in the training session using GNB, SVM, MLP, and LSTM decoders. **(c)** Cumulative fraction of the decoding error using different models. (One-way ANOVA, F(3,240)=45.02, p<0.0001; Bonferroni post-hoc test showed that decoding error using GNB was significantly higher than that using SVM, MLP, and LSTM, p<0.0001.) **(d)** Decoding performance using different models in the training session. The decoding model was constructed based on the first 75% of the recorded data and checked on the remaining 25%. The red curve demonstrates the mouse’s real position tracked by the video camera and the blue curve represents the reconstructed position. **(e)** The place field map of sensitive neurons in the real-time session. **(f)** The decoding error histogram in the real-time position reconstruction experiment. **(g)** An example of the mouse’s running trajectory reconstruction in a real-time session.

In the real-time session, the raw fluorescent intensities of location sensitive neurons detected in the training session were extracted and fed into the decoding model. We additionally plotted the place field map of location sensitive neurons in the real-time session, and all neurons showed clear specific firing locations (**Fig 3e**). We achieved a low average decoding error of 13.65 ± 0.50 cm/frame (13.20, 13.09, and 14.65 cm/frame, respectively) and a low median error of 9.33 ± 0.67cm/frame (8.00, 10.00, and 10.00 cm/frame). An example of the trajectory reconstruction is shown in **Fig 3f-g** and **Supplementary Fig 1a-b**.

We compared the noise levels of the signals in the training session and real-time session and did not detect significant differences (**Fig 4a-b**). The cross-correlation of place field maps between the two sessions showed a peak correlation value on the origin of the coordinate (**Fig 4c**), indicating stable neuronal firing patterns across sessions. We additionally tested the effects of LSTM window sizes. A 5-frame (∼0.17s) window size could provide an accurate decoding accuracy, while a 20-frame window size achieved much worse performance (**Fig 4d**). To test the effects of neuron numbers on the decoding accuracy, we compared the decoding accuracy using 20%, 40%, 60%, 80%, and 100% of sensitive neurons. Using more neurons in the reconstruction led to decreased decoding error (**Fig 4e**). Finally, we measured the processing time using different models and percent of neurons. The SVM model consumed considerably more time in processing each data frame (∼5ms using 100% sensitive neurons), while all the other models showed very fast processing speed (∼0.2ms using 100% sensitive neurons; **Fig 4f**).

**Figure 4.**
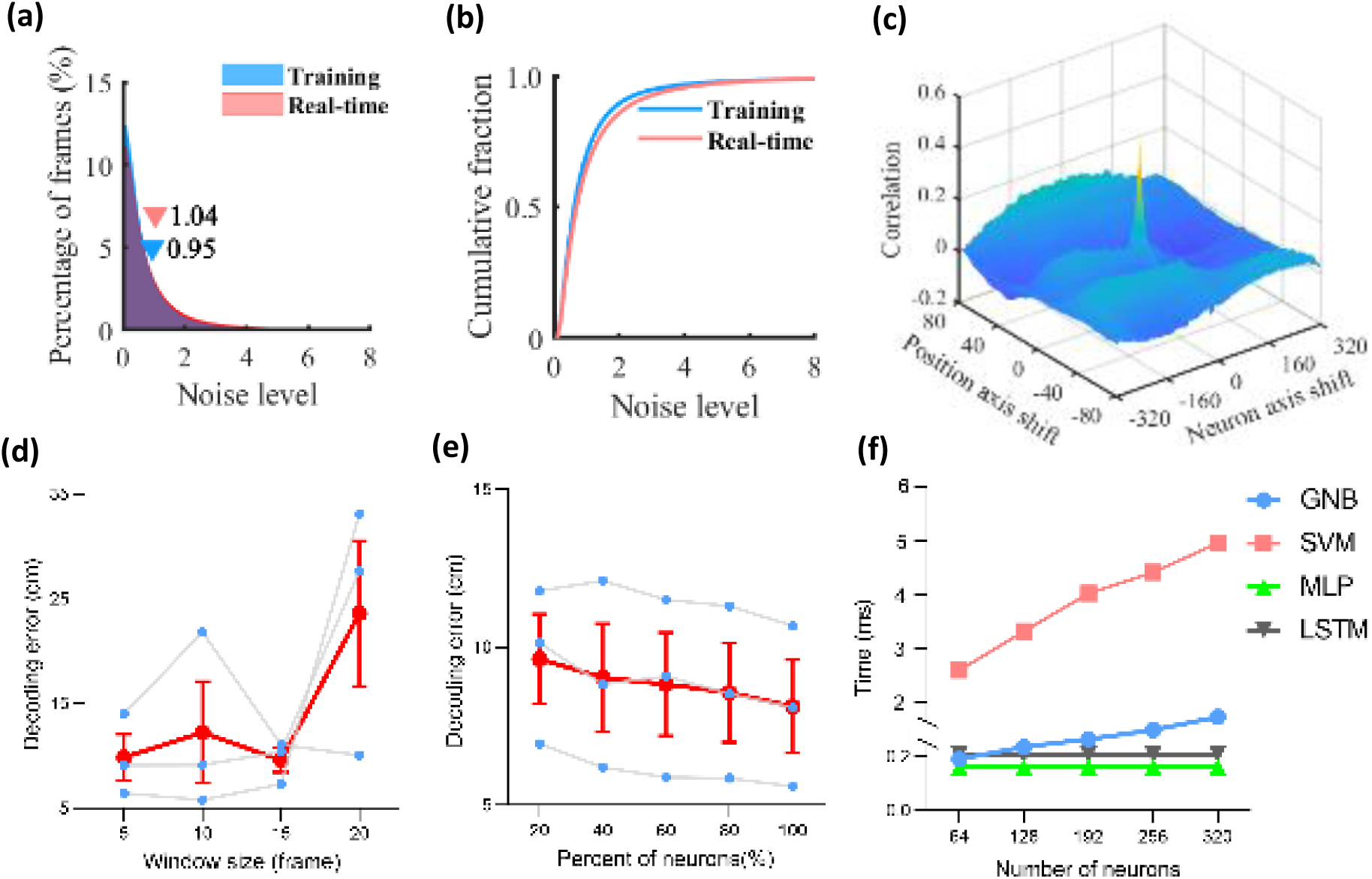
Position reconstruction decoding models. **(a-b)** Histogram and the cumulative fraction of the noise level in the training session and real-time session (paired Student’s t-test didn’t show significant differences: t=0.04121, df=79, p = 0.9672). **(c)** Cross-correlation of place field maps between the training session and real-time session. **(d)** Position reconstruction error as a function of time window size using the LSTM model. Blue dots represent the decoding error measured on three mice, and the red dots represent the mean error (Pearson correlation, *R*^2^ = 0.5646, p = 0.2486). **(e)** Position reconstruction error as a function of percent of sampled neurons using the MLP model. Error decreased by incorporating more neurons (Pearson correlation, *R*^2^ = 0.9716, p = 0.002). **(f)** Processing time as a function of the number of sampled neurons in a mouse using different models (Pearson correlation, GNB: *R*^2^ = 0.9926, p = 0.0003; SVM: *R*^2^ = 0.9876, p = 0.0006; MLP: *R*^2^ = 0.7701, p = 0.0505; LSTM: *R*^2^ = 0.5103, p = 0.1752).

### 3.2 Visual stimuli identification

We next studied whether the hippocampal neural ensemble activity could be decoded to identify visual inputs. A flashlight fixed on top of the recording chamber was switched on and off alternately, which was synchronized with the miniscope recording system. Three mice were tested individually in the experiment and each mouse first experienced a training session to construct the decoding model. There were 536, 707, and 569 neurons in the fields of view, respectively. The neuronal fluorescent intensity in light and dark environments showed similar distributions (**Supplementary Fig 2c**). The information content was then measured for each neuron, and we detected 204, 392, and 407 sensitive neurons in each mouse. An example of the average neuronal activity of sensitive neurons in light and dark epochs is shown in **Fig 5a**, while many neurons displayed strong responses during light-to-dark transient. We then tested the performance of a GNB decoder, an SVM decoder, and an MLP neural network to identify the visual inputs based on the training data. **Supplementary Table 2** summarizes the decoding accuracy of different models. The SVM decoder and MLP neural network both achieved very high decoding accuracy (mean decoding error: ∼3%), which was much higher than that using a GNB decoder (mean decoding error: 20.54% ± 8.70%). An example of the performance of different models is shown in **Fig 5b**. Again, different firing patterns of sensitive neurons were observed and were distinguishable in a high-dimensional neural state space (**Supplementary Fig 4.b,e)**. The model showing the best performance was applied in each mouse respectively in the real-time session.

**Figure 5.**
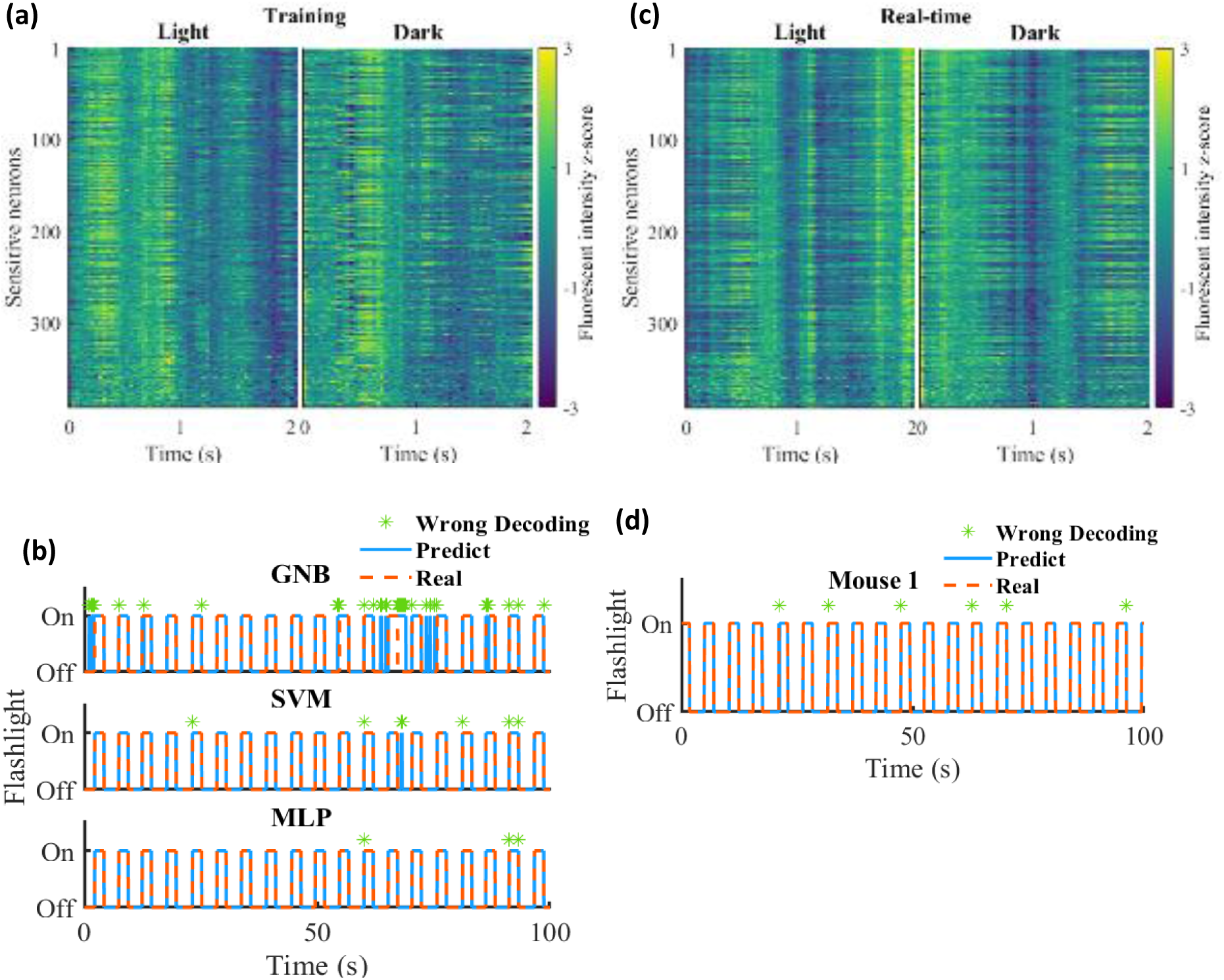
Visual stimuli experiment. **(a)** The visual stimuli evoked neuronal activity of sensitive neurons in light and dark epochs in the training session. The colour represents the fluorescent intensity z-score. The flashlight was turned on or off at time “0”. **(b)** An example of the decoding performance of different decoders in the training session (100*30 frames). The decoding model was constructed based on the first 75% data and checked on the remaining 25% data. The red dashed line and solid blue line represent the real and predicted status of the flashlight respectively. The wrong decoding frame is marked with a green star. **(c)** The visual stimuli evoked neuronal activity of sensitive neurons in light and dark epochs in the real-time session. **(d)** An example of the decoding performance in the real-time session (100*30 frames).

In the real-time session, the raw calcium signals of sensitive neurons were used to predict the visual inputs. Intriguingly, the average neuronal firing patterns looked different from those in the training session (**Fig 5c**) but decoding errors were very low in all three mice, which were 5.47%, 4.07%, and 0.53%, respectively (overall: 3.36% ± 1.47%). Examples of the decoding performance are shown in **Fig 5d** and **Supplementary Fig 2a-b**.

The noise levels of the signals in the training sessions and the real-time sessions did not show significant differences (**Fig 6a-b**). The maximum cross-correlation value of firing rate maps between the two sessions was not on the origin of the coordinate in this experiment, showing a shift along the time axis (**Fig 6c**). Considering more neurons resulted in higher decoding accuracy as observed in the position reconstruction experiment (**Fig 6d**). Because the decoding model was simpler than that in the position reconstruction experiment, the MLP model only required about 0.01ms to process the data in each frame (**Fig 6e**).

**Figure 6.**
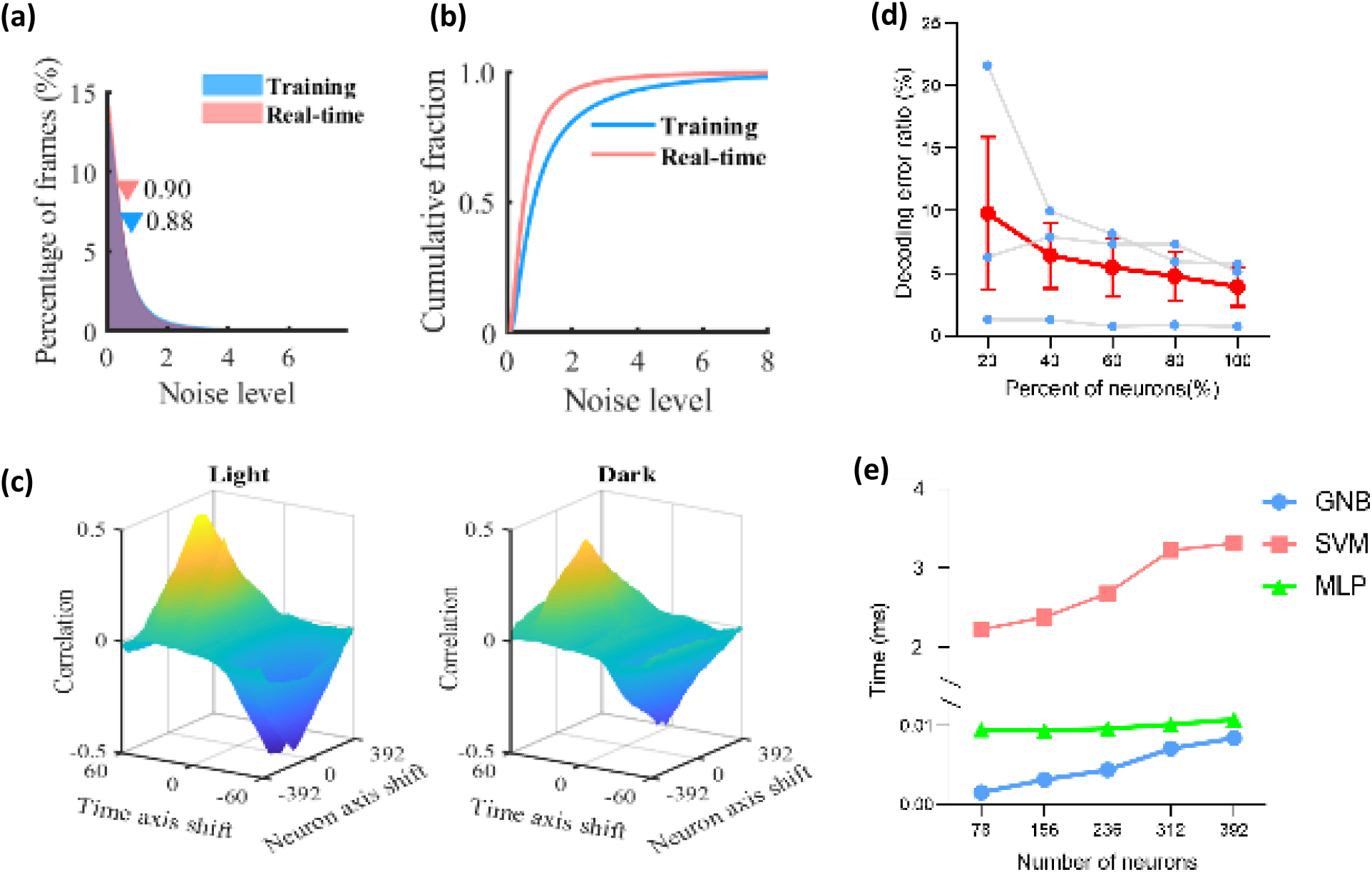
Visual stimuli decoding models. **(a-b)** Histogram and the cumulative fraction of the noise level in the training session and real-time session (paired Student’s t-test did not show significant differences: t=0.1362, df=79, p = 0.8920). **(c)** Cross-correlation of firing rate maps between the training session and real-time session in light and dark environments. **(d)** Decoding error as a function of percent of sampled neurons using the MLP model. Blue dots represent the decoding error measured on three mice, and the red dots represent the mean error. Error decreased considering more neurons (Pearson correlation, *R*^2^ = 0.8702, p = 0.0207). **(e)** Processing time as a function of the number of sampled neurons in a mouse using different models (Pearson correlation, GNB: *R*^2^ = 0.9853, p = 0.0008; SVM: *R*^2^ = 0.9523, p = 0.0045; MLP: *R*^2^ = 0.8162, p = 0.0355).

### 3.3 Auditory stimuli identification

To determine whether the hippocampal calcium activity could be decoded to distinguish the frequency of auditory stimuli, we exposed mice (n=3) to pure sinusoidal tones centred at 4kHz, 8kHz, or 16kHz. The speaker was activated and deactivated alternately, synchronizing with the miniscope recording system. In the training session, we observed 534, 619, and 599 neurons in each mouse, respectively. The animals experienced four different environments (mute, 4kHz, 8kHz, and 16kHz) in the experiment. We first attempted to use a GNB decoder, an SVM decoder, and an MLP neural network to identify the auditory input in each frame, but all the decoders failed to make accurate predictions. **Supplementary Table 3** summarizes the decoding error of each model, and several examples are shown in **Fig 7a**. We next sectioned the data into epochs according to the frequency of stimuli, and each epoch contained a 2s sound-on period followed by a 3s sound-off period. The neuronal fluorescent intensity in different audio frequency epochs showed similar distributions (**Supplementary Fig 3c**), but we detected 198, 227, and 230 sensitive neurons in each mouse, respectively. An example of the average neuronal activity of sensitive neurons in different environments is shown in **Fig 7b**. We then trained a CNN model to identify the frequency of each epoch. The CNN model achieved high decoding accuracy with an error rate of 17.67%, 22.76%, and 22.89% in each mouse, respectively (mean ± SEM: 21.11% ± 1.48%), based on the training data. An example of CNN performance is shown in **Fig 7c**. Again, different temporal firing patterns of sensitive neurons were observed, showing diverse features in a high-dimensional neural state space (**Supplementary Fig 4.c,f)**.

**Figure 7.**
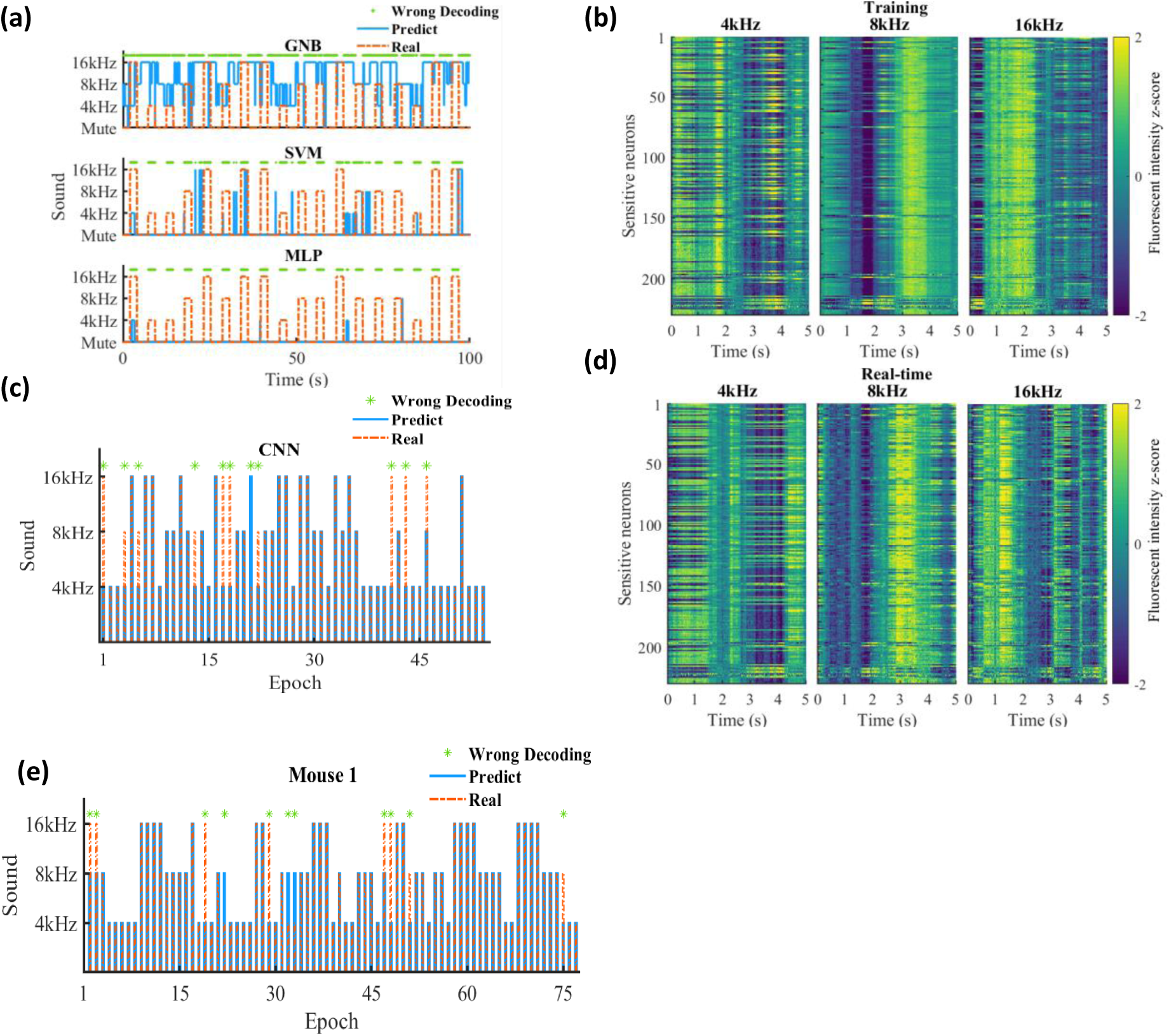
Auditory stimuli experiment. The mice were exposed to sound stimuli at three different frequencies. **(a)** An example of the decoding performance of different models in the training session (100*30 frames). The decoding model was constructed based on the first 75% data and checked on the remaining 25% data. The red dashed line and solid blue line represent the real and predicted status of the auditory stimuli respectively. The wrong decoding frame is marked with a green star. All decoders failed to make accurate predictions. **(b)** The sound stimuli evoked neuronal activity of sensitive neurons in 4kHz, 8kHz, and 16kHz epochs during the training session. The colour represents the fluorescent intensity z-score. The speaker was turned on at time “0” and turned off at time “2”. **(c)** An example of the CNN decoding performance in the training session. The data was sectioned into 5-s time-length epochs according to the frequency of the stimuli. **(d)** The sound stimuli evoked neuronal activity of sensitive neurons in the real-time session. **(e)** An example of the decoding performance using a CNN model in the real-time session.

In the real-time session, the CNN model constructed in the training session was deployed to decode the raw calcium activity of sensitive neurons. The average temporal firing patterns looked similar to those in the training session (**Fig 7d**). The decoding error ratio was 13.25%, 20.78% and 19.48% in each mouse, respectively (overall: 17.83% ± 2.32%). An example of the decoding performance is shown in **Fig 7e** and **Supplementary Fig 3a-b**.

The signals in the training and real-time sessions showed similar noise levels (**Fig 8a-b**). The cross-correlation of the firing rate maps between the two sessions showed several peaks with a dominant peak on the origin of the coordinate (**Fig 8c**), indicating relatively consistent temporal firing patterns across sessions. Likewise, using all sensitive neurons achieved the highest decoding accuracy (**Fig 8d**) and the model consumed about 1.6ms to process the 5s epoch data using a CNN model (**Fig 8e**).

**Figure 8.**
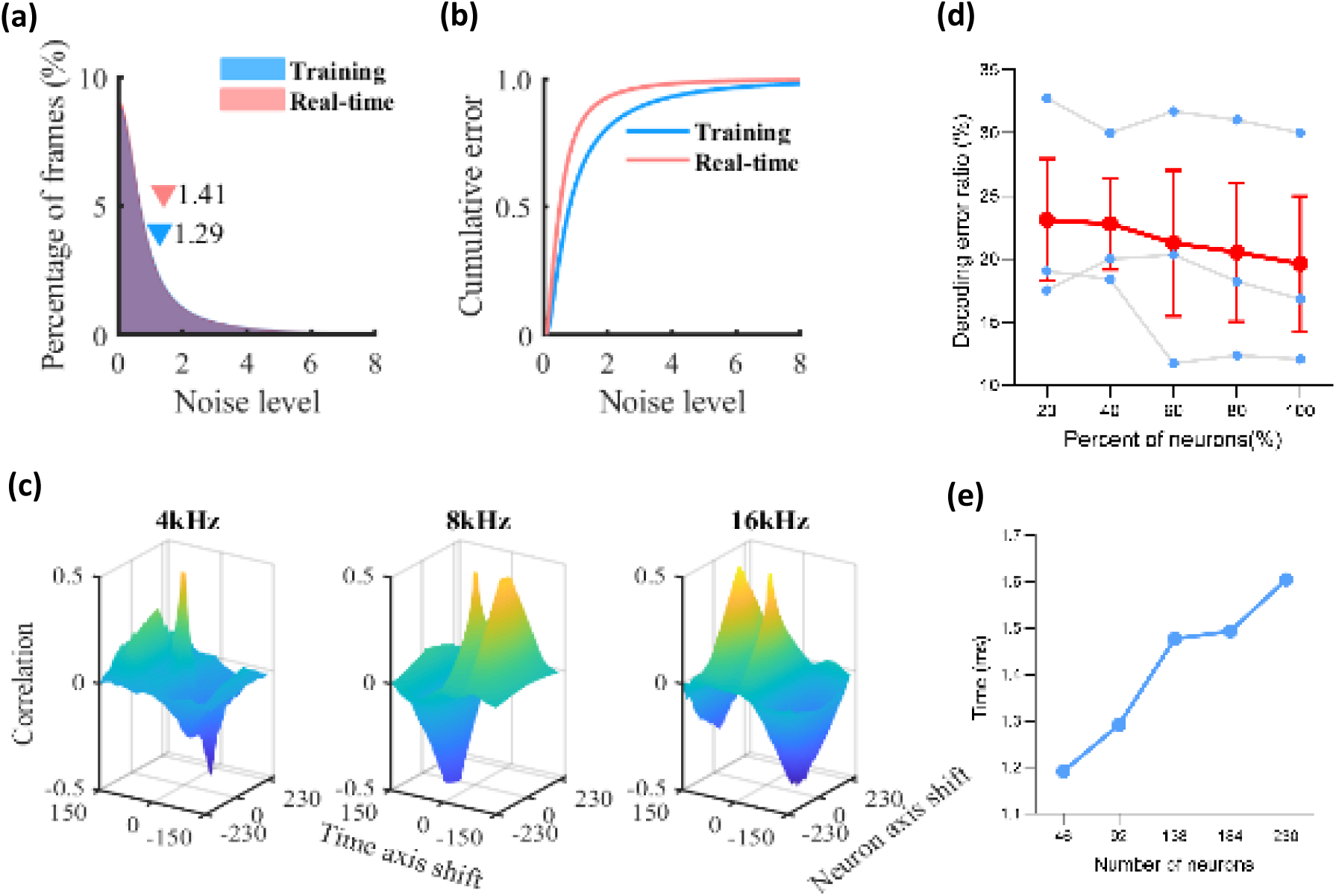
Auditory stimuli decoding models. **(a-b)** Histogram and the cumulative fraction of the noise level in the training session and real-time session (paired Student’s t-test did not show significant differences: t=1.226, df=79, p = 0.2238). **(c)** Cross-correlation of firing rate maps between the training session and real-time session in three environments. **(d)** Decoding error as a function of percent of sampled neurons using the CNN model. Blue dots represent the decoding error measured on three mice, and the red dots represent the mean error. Error decreased considering more neurons (Pearson correlation, *R*^2^ = 0.9717, p = 0.002). **(e)** Processing time as a function of the number of sampled neurons in a mouse using the CNN model (Pearson correlation, GNB: *R*^2^ = 0.9518, p = 0.0046;).

## Discussion

We have developed an optical brain-computer interface (OBCI) driven by calcium activity to decode separate sensory modality stimulation from hippocampal neuronal activity in mice. Our low-latency end-to-end analysis pipeline provides accurate decoding of both hippocampal spatial and non-spatial information, which facilitates direct neural communication.

### Hippocampal multi-sensory integration

The hippocampus supports a diverse cognitive map that incorporates both spatial and non-spatial information. Thus this structure may be a useful target to implement brain-computer interfaces to decode information from multi-sensory modalities, unlike other brain regions such as the visual cortex or auditory cortex that encode more specific sensory information.

The projection pathways of sensory information to the hippocampus differ anatomically. Spatial information mainly targets the dorsal and posterior hippocampus (Strange et al., 2014). Previous work using electrophysiological data was able to extract spatial information from the hippocampus in real time with a small set of neurons. Guger et al. (2011), Sodkomkham et al. (2016), and Hu et al. (2018) reconstructed the running trajectory of rats in real time by recording hippocampal action potentials.

Non-spatial information mostly flows into the ventral and anterior hippocampus (Strange et al., 2014). The visual signal projects to the hippocampus from the visual cortex through a multi-synaptic pathway (Lavenex and Amaral, 2000; Ranganath and Ritchey, 2012). Additionally, Haggerty and Ji (2015) observed the synchrony between activity of hippocampal neurons and visual cortical neurons in freely moving rats. The transmission pathways between the hippocampus and auditory cortex are more complex. It is believed that there are two major pathways – the lemniscal pathway and the non-lemniscal pathway, and the auditory signalling to hippocampus has likely undergone several integrative stages (Munoz-Lopez et al., 2010; Xiao et al., 2018). These studies demonstrate the complex connectivity basis of highly processed multi-sensory encoding in the hippocampus and provide insight into the difficulty of decoding these integrated signals.

### Decoding models

We tested different machine learning models to decode the hippocampal cognitive map including a GNB model, an SVM model, an MLP model, an LSTM model, and a CNN model. In all experiments, the GNB model showed the highest decoding error, which might be due to the nonlinearity of the hippocampal neuronal network and that the cognitive map was described in complex high-dimensional spaces. But on the other hand, these results are not surprising, because it was not straightforward to determine appropriate priors for the GNB model.

Intriguingly, in both position reconstruction and visual stimuli identification experiments, MLP and SVM models showed similar decoding performance in all animals. Compared with an MLP model, an SVM model has the advantage of fewer hyper-parameters to optimize. Additionally, an SVM model can be trained in an online mode (Jain et al., 2014; Laskov et al., 2006), which avoids a separate training session.

The long short-term memory (LSTM) algorithm is an artificial recurrent neural network that can achieve good prediction accuracy from time-series data (Rezaei et al., 2018; Tampuu et al., 2019). It simulates a biologically relevant model of how neuronal activity is processed. However, it did not show the best performance in the position reconstruction experiment, which was somewhat surprising. A possible explanation may be the slow kinetics of neuronal calcium activity. Action potentials cause calcium influx and efflux in excitable cell bodies, and the depolarization-evoked neuronal firing has a long-lasting effect on calcium activity. This indicates that the calcium activity in the current frame inherits partial information encoded in previous frames, which as a result weakens the strength of memory units in the model. Intriguingly, incorporating too much history (very old information) in an LSTM network causes a drop in its performance because that information may not be relevant or useful or may introduce unwanted noise.

Spatial and visual information could be decoded accurately in each frame in the experiments. However, a relatively long-time window was needed to decode the auditory information. This may be due to the different encoding mechanisms of hippocampal neurons for different types of sensory information. Auditory-evoked neuronal activation has been reported to exhibit variable latencies (MacDonald et al., 2011; Itskov et al., 2012), resulting in long-lasting temporal firing patterns. Another explanation may be the relatively low absolute sensitivity of CA1 neurons to auditory stimuli (Moita et al., 2003; Itskov et al., 2012), necessitating the highly optimised signal analysis techniques needed for this task.

### Optical Brain-Computer Interfaces (OBCIs)

Conventional intracranial brain-computer interfaces (BCIs) use electrode arrays to record neuronal action potentials or local field potentials. More recently, the potential for calcium imaging using multiphoton of neuronal ensembles as a BCI has been examined (Clancy et al., 2014; Trautmann et al. 2021). The relationship between neuron action potentials and calcium activity is complex. An action potential activates voltage-gated calcium channels, eliciting a nonlinear rise in intracellular calcium concentration. Although the dynamics of calcium activity is relatively slow, it has been shown to track action potential frequency (Harding et al., 2020). In our experiments, we used GCaMP6f with relatively fast kinetics (∼50ms temporal resolution; Chen et al., 2013). Thus, the temporal resolution of calcium signals seems functionally comparable to electrical signals.

Clancy et al. (2014) utilized volitional control of a small number of neurons identified with two-photon imaging that could indirectly control the sound from a speaker. Trautmann et al. (2021) implemented a two-photon OBCI in a head-fixed macaque for detection of the animal’s arm motion. In contrast, our technique uses single-photon imaging, which in practice can detect many more neurons, allowing directly decoding of spatial, light and sound modalities at high precision in real time. These remarkable capabilities derive from the use of a large neural dataset, allowing the use of machine learning algorithms for very low latency signal processing. Additionally, this workflow is low-cost and does not require bulky equipment.

In summary, we constructed an OBCI system and successfully decoded spatial, visual, and auditory information from the mouse hippocampus. The end-to-end OBCI system proposed here presents a proof of concept for decoding the hippocampal cognitive map in real time. It expands the method and opportunity to study the activity of hippocampal neuronal ensembles and will be helpful for future content-specific closed-loop BCI experiments. Furthermore, it provides an approach for “cognitive decoding”, which may be applied in clinical applications and scientific research in the future.

## Author Contributions

D.S., R.R.U and C.F. conceived and designed the study. D.S. carried out the experiment, processed the data, and drafted the manuscript. F.H. and Y.Y processed the data. R.R.U and C.F supervised the project and did critical manuscript revision. All authors have read and approved the final manuscript.

## Conflict of Interest

The authors declare that the research was conducted in the absence of any commercial or financial relationships that could be construed as a potential conflict of interest.

## Data availability statement

All data supporting the findings of this study are available from the corresponding authors upon reasonable request.

## Funding

This work was supported by Royal Melbourne Hospital Neuroscience Foundation (A2087) and Australian Research Council under Discovery Project (DP170100363).

---

**Supplementary Figure 1.**
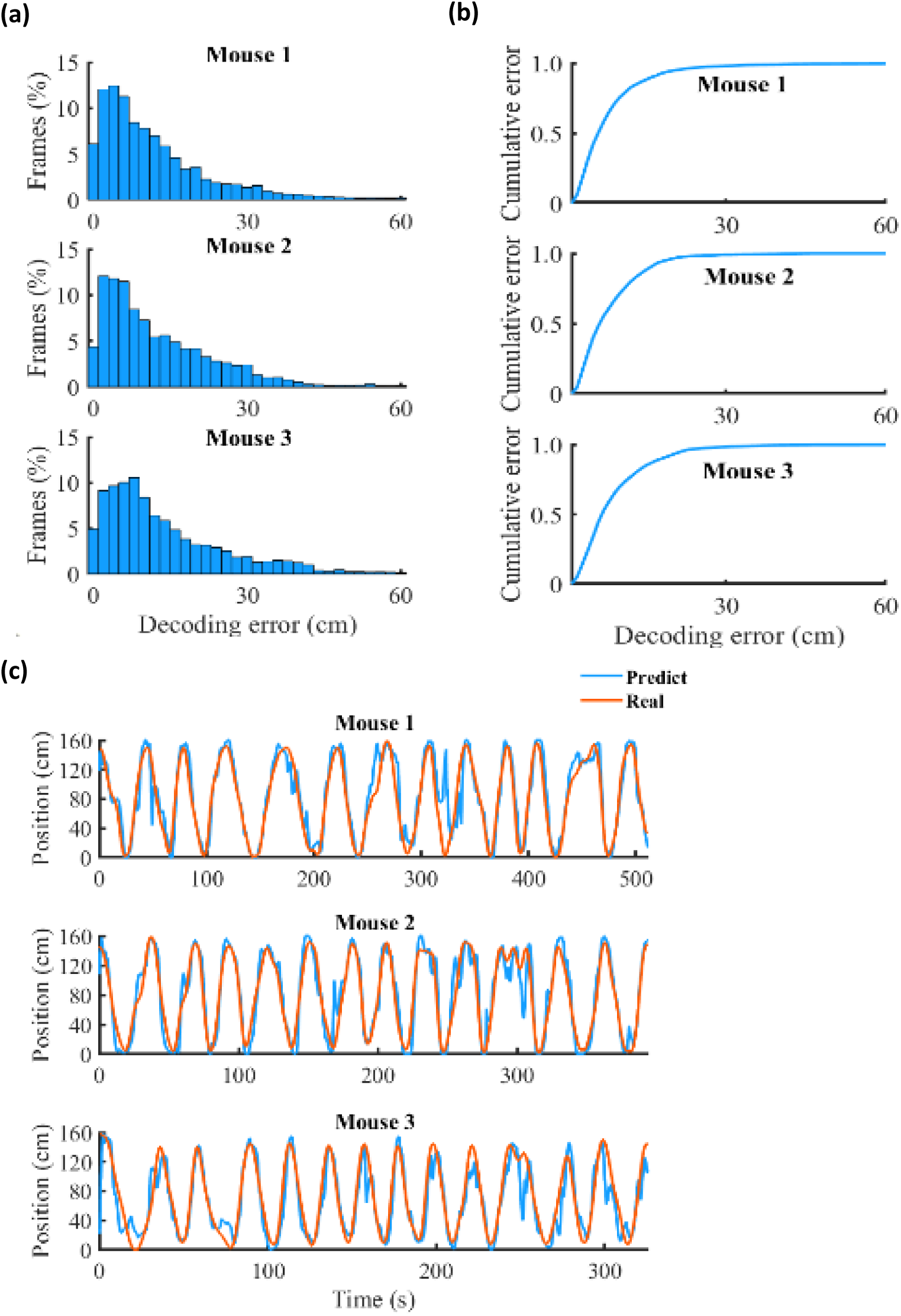
Position reconstruction experiment. **(a-b)** Histogram and the cumulative fraction of the decoding error in the real-time session. The decoding model with the best performance in the training session was implemented in each mouse respectively. **(c)** Mice’s running trajectory reconstruction in the real-time session. The red curve demonstrates the mouse’s real position tracked by the video camera and the blue curve represents the reconstructed position.

**Supplementary Figure 2.**
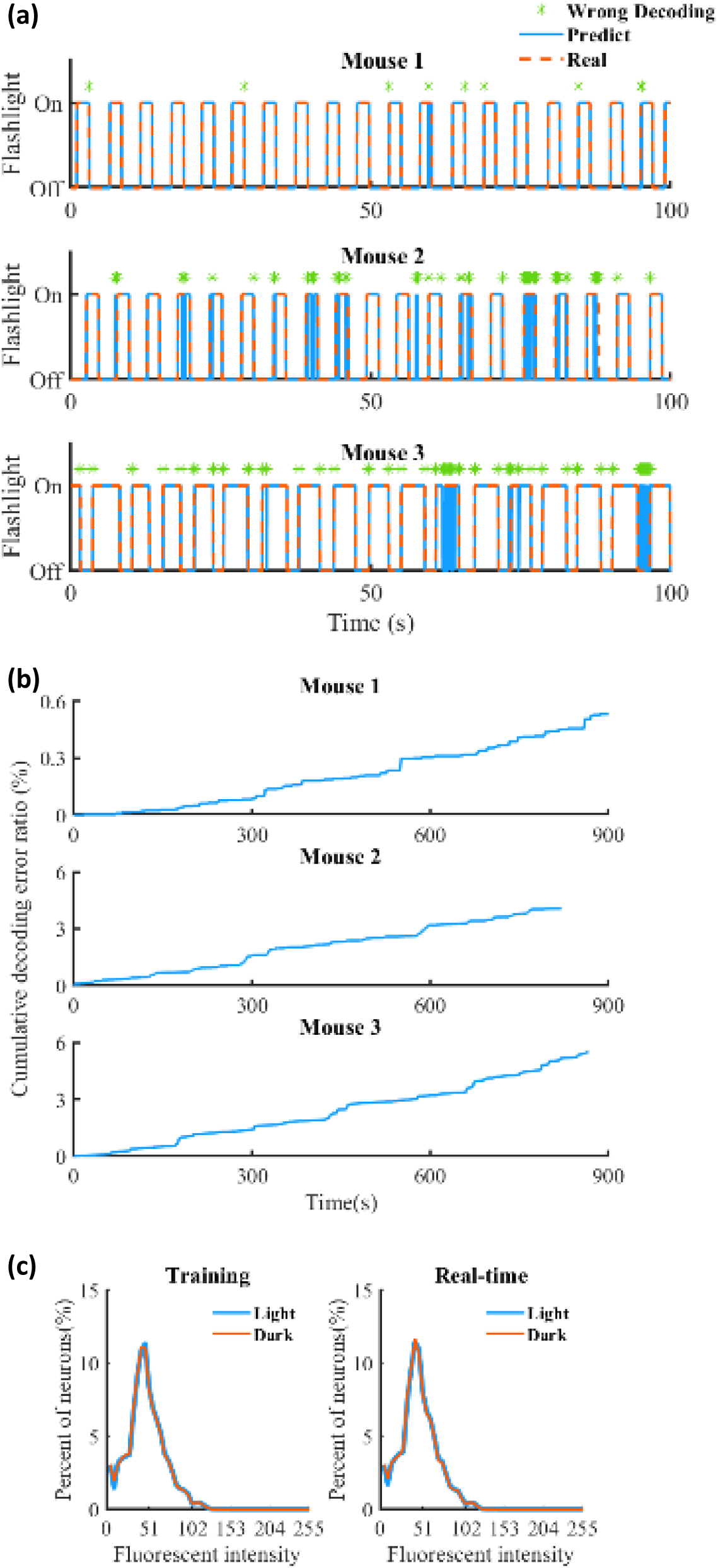
Visual stimuli identification experiment. **(a)** Examples of the decoding performance in the real-time session (100*30 frames). The decoder with the best performance in the training session was used in each mouse respectively. The red dashed line and solid blue line represent the real and predicted status of the auditory stimuli respectively. The wrong decoding frame is marked with a green star. (b) Cumulative decoding error ratio with respect to time. **(c)** Distributions of neurons’ fluorescent intensity. Paired Student’s t-test didn’t show significant differences (Training: t= 4.403×10^−14^, df=51, p > 0.9999; Real-time: t=9.703×10^−14^, df=51, p > 0.9999).

**Supplementary Figure 3.**
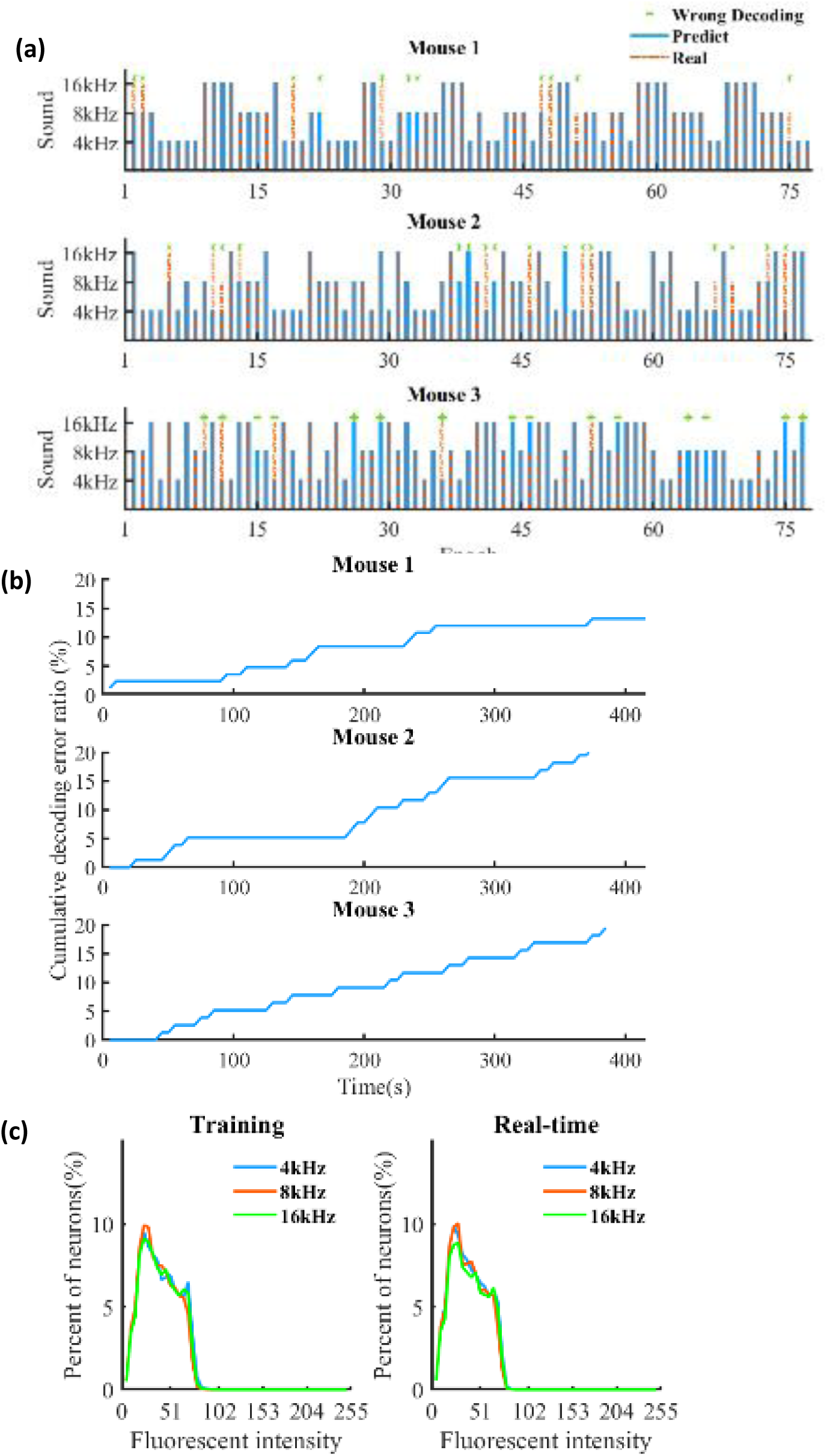
Auditory stimuli identification experiment. **(a)** Examples of the decoding performance using a CNN model in the real-time session. The red dashed line and solid blue line represent the real and predicted status of the auditory stimuli respectively. The wrong decoding frame is marked with a green star. (b) Cumulative decoding error ratio with respect to time. **(c)** Distributions of neuronal fluorescent intensity. One-way ANOVA didn’t show significant differences (Training: F(2,102)=0.4342, p =0.6490; Real-time: F(2,102)=2.853, p =0.0623).

**Supplementary Figure 4.**
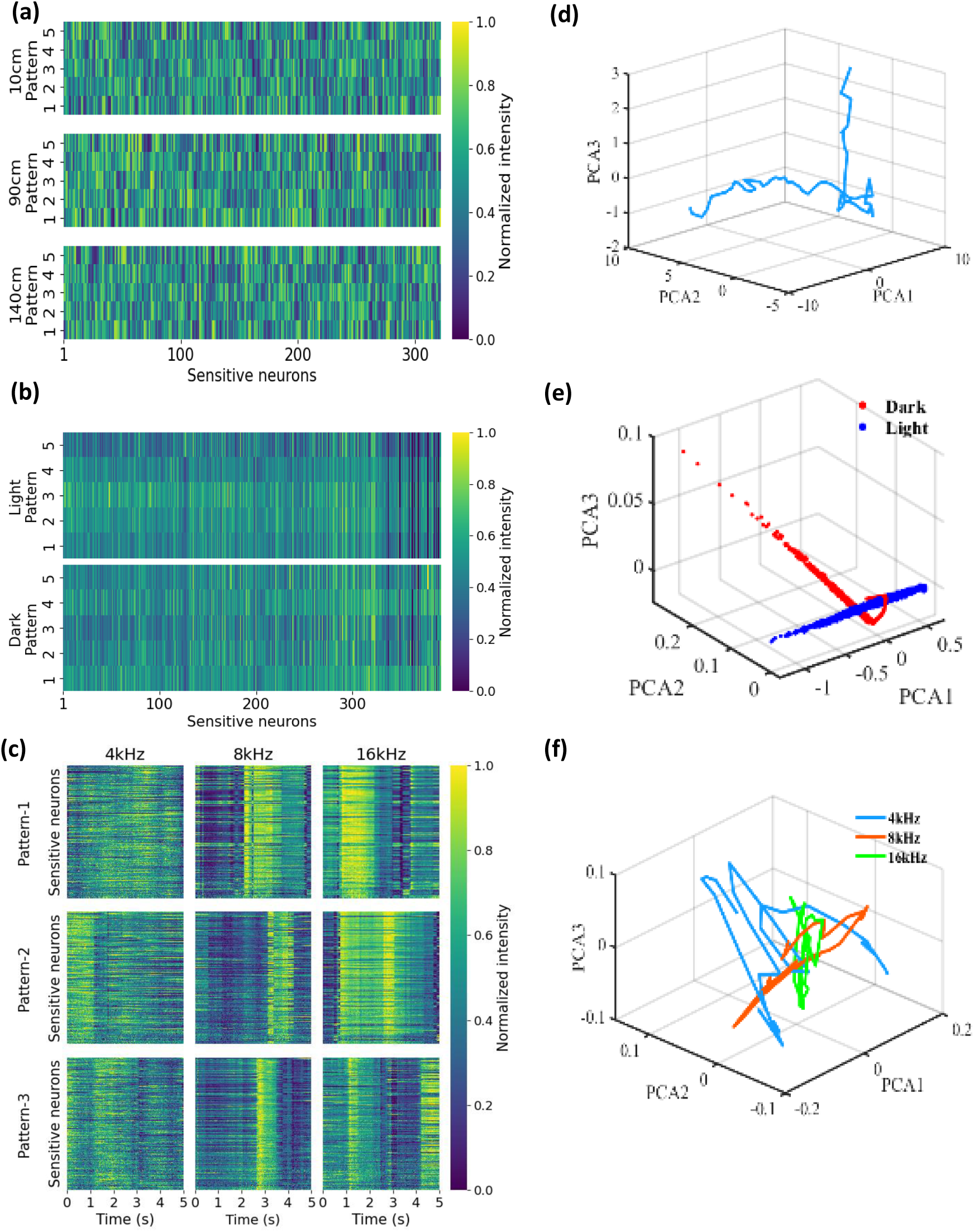
Neuronal ensemble firing patterns. Some examples of sensitive neurons’ firing patterns in **(a)** linear track experiments, **(b)** visual stimuli experiments, and **(c)** auditory stimuli experiments. **(d-f)** show the animal’s running trajectory, visual stimuli responses, and auditory stimuli responses described in a high-dimensional neural state space. Features were extracted from the output of the last hidden layer in a MLP model **(d-e)** or a CNN model **(f)**. Principle component analysis (PCA) were applied on the features for better visualization.

**Supplementary Table 1.** The mean and median decoding error (cm/frame) in the training session of the position reconstruction experiment. The first row in each cell represents the mean error and the second row represents the median error. The overall error is expressed as mean/median ± standard error mean (SEM).

| Decoder / mean<br>median | Mouse_1 | Mouse_2 | Mouse_3 | All (cm/frame) |
| --- | --- | --- | --- | --- |
| **GNB** | 17.76<br>12 | 20.00<br>14 | 30.60<br>22 | 22.79 ± 3.42<br>16.00 ± 2.64 |
| **SVM** | 10.45<br>8 | 16.42<br>12 | 13.06<br>10 | 13.31 ± 1.49<br>10.00 ± 1.00 |
| **MLP** | 10.29<br>8 | 16.24<br>12 | 16.98<br>14 | 14.50 ± 1.83<br>11.33 ± 1.52 |
| **LSTM** | 11.16<br>10 | 17.13<br>12 | 16.44<br>14 | 14.91 ± 1.63<br>12.00 ± 1.00 |

**Supplementary Table 2.** The average decoding error ratio in the training session of the light stimuli experiment. The overall error ratio is expressed as mean ± SEM.

| Decoder | Mouse_1 | Mouse_2 | Mouse_3 | All |
| --- | --- | --- | --- | --- |
| **GNB** | 22.95% | 34.28% | 4.40% | 20.54% ± 8.70% |
| **SVM** | 3.99% | 4.01% | 1.22% | 3.08% ± 0.92% |
| **MLP** | 5.36% | 4.63% | 0.73% | 3.57% ± 1.43% |

**Supplementary Table 3.** The average decoding error ratio in the training session of the sound stimuli experiment. The overall error ratio is expressed as mean ± SEM.

| Decoder | Mouse_1 | Mouse_2 | Mouse_3 | All |
| --- | --- | --- | --- | --- |
| **GNB** | 71.09% | 77.50% | 81.34% | 76.64% ± 2.59% |
| **SVM** | 42.81% | 39.85% | 40.05% | 40.91% ± 0.82% |
| **MLP** | 39.33% | 37.16% | 37.84% | 38.11% ± 0.55% |
| **CNN** | 17.67% | 22.76% | 22.89% | 21.11% ± 1.48% |

## Notes

### Competing Interest Statement

The authors have declared no competing interest.

